# The Contribution of the Koniocellular Visual Pathway to Aversive Learning in Human Visual Cortex

**DOI:** 10.1101/2025.04.24.650318

**Authors:** Katherine J. McCain, Nathan M. Petro, Andreas Keil

## Abstract

The present study examined the contribution of the koniocellular retino-geniculate visual pathway to the electrocortical amplification of threat cues in human visual cortex using an aversive conditioning task. The task involved tritan stimuli, which are thought to convey visual information through S-cone signals that project to regions along the koniocellular pathway (tritan condition) and achromatic stimuli that preferentially activate luminance channels (luminance condition). Steady-state visual evoked potential (ssVEPs) responses to the conditioned threat (CS+) and safety cues (CS-) in each condition were analyzed using a non-parametric Bayesian bootstrapped approach. Results showed that the tritan and luminance conditions exhibited greater ssVEP responses to the CS+ compared to the CS-stimuli in occipital sensors early into the trial (0 ms - 1000 ms; logBF10 > 2, decisive support). In addition to these early conditioning effects, a late conditioning effect was observed (1500 ms - 2500 ms) in the tritan condition that emerged in bilateral anterior sensors (logBF10 > 2). To further examine the tritan contribution to aversive learning, transitive Bayes factors were computed to compare the magnitude of the conditioning effects across conditions. Transitive Bayes factors showed that the early conditioning effect was larger in the luminance condition compared to the tritan condition (logBF10 > 2). Furthermore, the late conditioning effect remained larger in the tritan condition compared to the luminance condition (logBF10 > 2). Our findings suggest that both the koniocellular visual pathway and luminance channels play a role in the electrocortical amplification of threat signals in human visual cortex.

---

Interactions between local inhibitory circuits in primary visual cortex (V1) and higher-order feedback projections are thought to contribute to the sharpening of excitatory lateral geniculate nucleus (LGN) afferents, facilitating the selective visuocortical amplification of motivationally relevant information (Angelucci et al., 2002; Keil et al., 2007; Briggs, 2020; Li and Keil, 2023). Several, electrophysiological studies have demonstrated that (Song and Keil, 2014; McTeague et al., 2015; Friedl and Keil, 2021) aversive conditioning resulted in heightened visuocortical responses to conditioned threat cues. These studies conveyed visual threat through luminance channels which primarily involve magnocellular and parvocellular retino-geniculate signals (Vidyasagar et al., 2002; Johnson et al., 2004). However, how the koniocellular retino-geniculate pathway contributes to the acquisition of aversive contingencies has yet to be established. Therefore, the present study aimed to investigate the koniocellular contribution to aversive learning in human visual cortex with a simple aversive conditioning paradigm.

In primates, neurons in the koniocellular layers of the LGN receive blue-on yellow-off color signals from bistratified retinal ganglion cells and transmit these signals to the cytochrome oxidase blobs in layer 2/3 of V1 (Livingstone and Hubel, 1984; Hendry and Yoshioka, 1994; Martin et al., 1997; Hendry and Reid, 2000; Dacey, 2000; Casagrande et al., 2007). Blue-on yellow-off signals can arise from short-wavelength cone (S-cone) stimulation while adapting the medium-(M) and long-wavelength (L) cones using the tritanopic technique proposed by Cavanagh and colleagues (1992). Furthermore, Tritan stimuli are near-equiluminous and attenuate minor S-cone contributions to luminance channels (Lee and Stromeyer, 1989; Chatterjee and Callaway, 2002). Several studies have utilized S-cone biasing stimuli and observed electrocortical amplification of the S-cone opponent signals in humans and primates (Cottaris and De Valois, 1998; Mullen et al., 2007, 2008). For instance, Mullen and colleagues (2008) identified a differential increase in BOLD activation in response to S-cone biasing stimuli in humans: S-cone biasing stimuli prompted small BOLD responses at the level of the LGN, but strong responses in V1. Therefore, based on this evidence, it is plausible that the koniocellular pathway is involved in the visuocortical amplification of S-cone opponent signals, and that the enhancement of this signal may occur at the cortical level.

Additional support for the notion that the koniocellular pathway plays a role in the modulation of visuocortical activation is derived from evidence of neurophysiological connections to subcortical and extrastriate regions in primates that are thought to be involved in the detection and evaluation of salient visual information (Casagrande, 1994; Hendry and Reid, 2000; Sincich et al., 2004; Isbell, 2006; Nassi and Callaway, 2009). The superior colliculus, involved in early encoding of salient visual information and the execution of ocular movements (Sprague and Meikle, 1965; Krebs et al., 2010; White et al., 2017), exhibits afferents that project to the koniocellular layers of the LGN in primates (Casagrande, 1994; Hendry and Reid, 2000). In addition, the koniocellular layers of the LGN exhibit a direct projection to the motion sensitive middle temporal (MT) region of extrastriate cortex in primates (Sincich et al., 2004). Findings in humans suggest that the koniocellular pathway is involved in fast saccadic responses to complex biologically relevant S-cone biasing stimuli (Kveraga et al., 2020). Similarly, it has been established that the koniocellular pathway is sensitive to motion and exhibits visuocortical activation to moving S-cone biasing stimuli (Wandell et al., 1999; Morand et al., 2000).

In the present study, we leveraged the properties of steady-state visual evoked potentials (ssVEPs) to measure selective visuocortical responses to conditioned threat (CS+) and safety cues (CS-). S-cone biasing tritan stimuli were used to convey visual information to the koniocellular visual pathway, and achromatic luminance stimuli were used to bias luminance channel responses. We expected the tritan stimuli to elicit robust electrocortical responses, in line with several studies that observed substantial electrocortical activity in response to S-cone biasing stimuli (Cottaris and De Valois, 1998; Mullen et al., 2007, 2008). Furthermore, we expected greater ssVEP responses to the CS+ stimuli, compared to the CS-stimuli in both the tritan and luminance conditions (Song and Keil, 2014; McTeague et al., 2015; Li and Keil, 2023). Finally, we measured pupil dilation as a manipulation check, expecting greater re-dilation for the CS+ compared to the CS-stimuli in each condition because pupil re-dilation is an index of emotional arousal and sympathetic activation (Bradley et al., 2008).

## Methods

### Participants

Fifty (mean ± SD, 19.46 ± 1.95 years; 36 female) undergraduate students were recruited at the University of Florida in exchange for course credit. All participants provided written informed consent and reported normal or corrected-to-normal vision and a negative personal and family history of seizures. All study procedures were in accordance with the Declaration of Helsinki, and approved by the University of Florida Institutional Review Board.

### Experimental design

#### Stimuli

Visual stimuli were developed in Matlab (MathWorks) using Psychophysics Toolbox version 3.0.15 (Brainard, 1997) and presented on a Cambridge Research Systems Display ++ monitor (1,920 × 1,080 pixels, 120 Hz refresh rate) connected to a PC running Linux Ubuntu. Each trial began with an adapter pre-stimulus period (3-7 seconds) that consisted of a fixation dot (visual angle 0.19°) in the center of a bright yellow background (190, 190, 0 RBG; 55.30 cd/m^2^) in tritan conditions or a black background (0, 0, 0 RBG; 0.05 cd/m^2^) in luminance conditions. Following the adapter pre-stimulus period, the oriented (45° or 135° clockwise tilt) flickering gratings were presented for 4 seconds. Tritan gratings were created by adding the blue RGB channel (190, 190, 230 RBG; 58.86 cd/m^2^) to defined alternating regions of a central aperture in the yellow adapter (see figure 1). The resulting stimulus appeared to observers as a white and yellow grating with a visual angle of 3.10° and was flickered at 7.5 Hz (driving frequency), exhibiting a spatial frequency of 16 cycles/°. Luminance gratings were created by adding white (255, 255, 255 RBG; 78.95 cd/m^2^) to defined alternating regions of a central aperture in the black adapter and exhibited the same visual angle, flicker frequency, and spatial frequency as the tritan gratings (see figure 1). The bright yellow adapter pre-stimulus period functions to minimize the luminance sensitivity of M- and L-cones, so that slight changes in luminance (tritan grating presentation) do not surpass their luminance contrast threshold. As a result, the tritan gratings appear near-equiluminous, and the opponent signals conveyed by the tritan grating should predominantly be transmitted through S-cones (Cavanagh et al., 1992). The Michelson contrast of the tritan gratings was 0.03 and 0.99 for the luminance gratings. Visual stimuli were measured using a Gossen MavoSpot Luminance meter.

**Figure 1.**
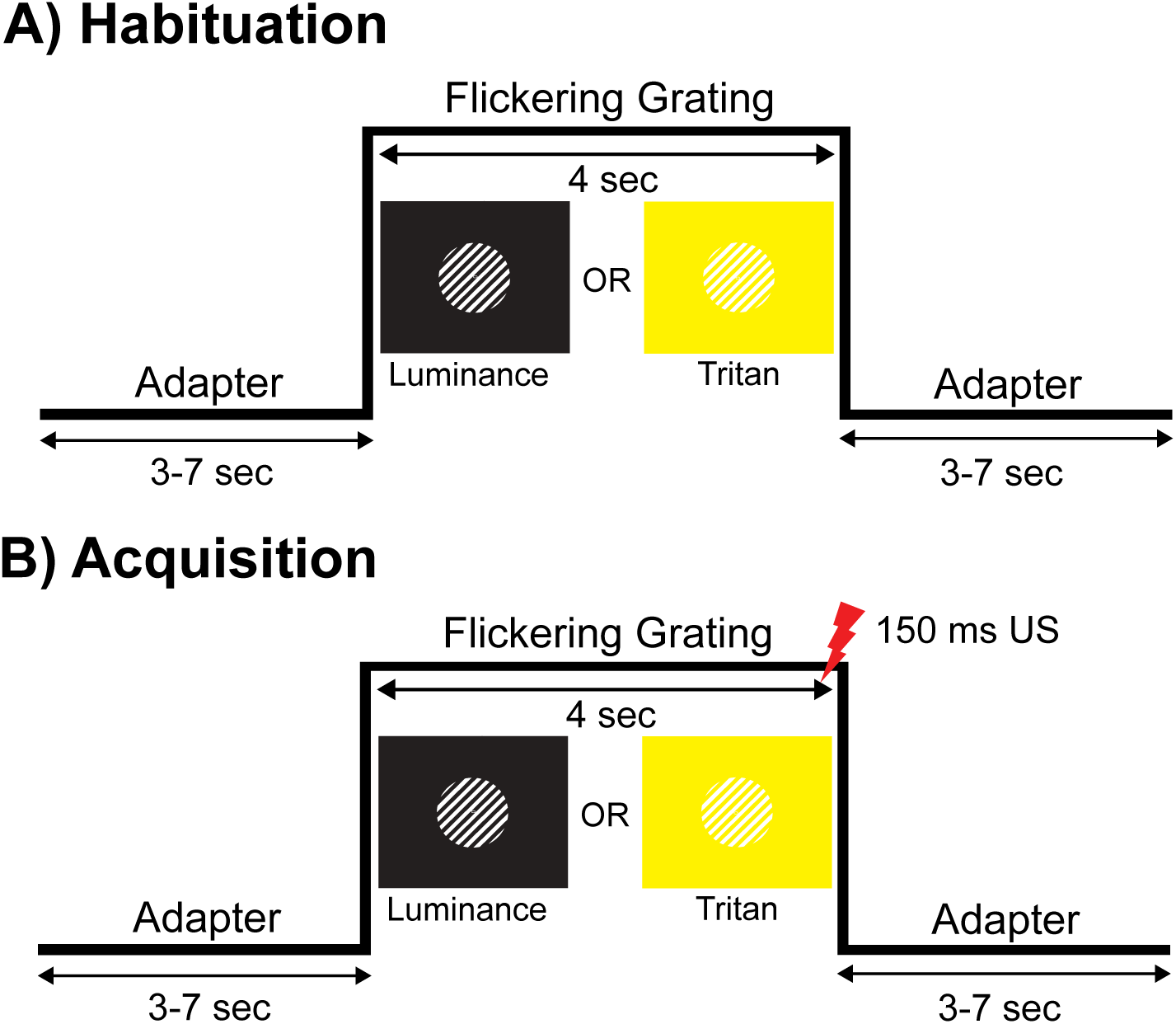
Experimental paradigm. The aversive conditioning task consisted of luminance and tritan conditions. Trials involved adapter periods (3-7 sec) that flanked the flickering grating (7.5 Hz) period (4 sec). The adapter preceding the flickering grating was always a black background in luminance conditions and a bright yellow background in tritan conditions. (A) In the habituation phase, the flickering grating was presented without any US (mild electrical stimulus) pairings. (B) In the acquisition phase, the orientation (45° or 135°) of the flickering grating signaled the presence (CS+) or absence (CS-) of the US (150 ms) and was counterbalanced across participants.

The unconditioned stimulus (US) was a mild electrical pulse administered using a Digitimer DS7A constant current stimulator (Hertfordshire, United Kingdom) connected to an electrode placed on the participant’s non-dominant wrist. The pulse duration was 150 ms and co-terminated with the flicking grating in trials where the orientation of the grating was paired with the mild electrical stimulus.

#### Procedure

Participants were seated about 120 cm in front of the stimulus presentation monitor in an electrically shielded, sound-attenuated room with dim lighting. Once the EEG sensor net was applied, the eye-tracking system and electrical stimulus intensity were adjusted and individually set for each participant. Before beginning the experiment, participants were provided written and verbal instructions describing that they would see a series of flickering patterns on the screen and would occasionally feel an annoying electrical stimulus. Participants completed expectancy ratings throughout the task and were asked to rate the likelihood that they would expect to feel the annoying electrical stimulus when shown each flickering pattern. The expectancy ratings involved dragging the mouse pointer across a scale where the far left of the scale indicated “not likely that the flickering pattern was accompanied by a shock”, the middle indicated “uncertain”, and the far right indicated “certain that the flickering pattern was accompanied by a shock”.

The intensity of the electrical stimulus was determined during the shock work-up phase and was set individually for each participant prior to beginning the task. The shock work-up consisted of repeated electrical stimulus pulses that started at 0 mA and increased incrementally by 0.1 mA. After each electrical stimulus pulse, the participants rated the intensity of the stimulus using a pain scale where 0 indicated that the stimulus was not annoying, aversive, or painful and 10 indicated that the stimulus was extremely painful. The intensity was increased until the participants rated the electrical stimulus as a level 4 on the pain scale, indicating that the stimulus was highly aversive and uncomfortable, but not painful. The shock work-up consisted of two separate work-ups to a level 4 on the pain scale. The intensity level was recorded after each work-up, averaged together, and then an additional 25% of the average intensity level was added to reach the final intensity used throughout the experiment. The final intensity level was delivered to the participant immediately before beginning the experiment to ensure that it was still tolerable and not painful. If the final intensity was reported to be painful and no longer tolerable, the intensity was reduced to the mean amperage prior to the 25% increase. However, if the intensity was still intolerable then the intensity was reduced until it was tolerable for that participant. The average electrical stimulus intensity was 0.87 ± 0.68 mA (M ± SD).

The task consisted of 160 trials, with 20 trials in the habituation and extinction phases and 120 trials in the acquisition phase. During the habituation phase, participants were introduced to each of the stimulus types without the presence of the mild electrical stimulus (US) and prompted to provide expectancy ratings after completing the 20 trials. In the acquisition phase, the US was introduced, and the orientation (45° or 135° clockwise tilt) of the grating signaled the presence (CS+) or absence (CS-) of the mild electrical stimulus in the tritan and luminance conditions. Participants provided expectancy ratings after the first 20 trials of the acquisition phase and then again after 80 trials. The orientation of the CS+ grating was counterbalanced across participants and the reinforcement rate during the acquisition phase was 100%. The extinction phase was included primarily for ethical reasons to reduce the association between the CS+ and the US. Participants also completed expectancy ratings after the 20 trials of extinction. Throughout the task, the occurrence of a given trial type was equiprobable and pseudo-randomized.

### Data acquisition and processing

#### Eye-tracking

Pupil data was recorded with an EyeLink 1000 Plus system on a 16mm lens (SR Research) and sampled at 500 Hz. The data was filtered offline using a Butterworth 6^th^ order low-pass filter with a cutoff frequency of 0.03 Hz. Next segments of 600 ms pre-stimulus onset and 3800 ms post-stimulus onset were extracted from the continuous data, and segments with missing pupil data were identified and estimated using cubic interpolation. The resulting segments were averaged across conditions and baseline corrected to the average diameter of the 200 ms prior to stimulus onset.

#### EEG

Continuous electroencephalogram (EEG) data was recorded using a 129-electrode high-density EEG System (EGI) with Cz as the recording reference and sampled at 500 Hz. Offline data processing was completed using a custom automated EEGLAB pipeline in Matlab (Delorme and Makeig, 2004), and the code can be found here https://osf.io/yue78/. Continuous EEG data was filtered using a 3 to 25 Hz Butterworth 4^th^ order bandpass filter, and eye blinks were corrected using the automated eye correction procedure developed by Schlögl et al. (2007) using the BioSig open-source package for biomedical signal processing (BioSig, 2005). Next, epochs of 600 ms pre-stimulus to 3800 ms post-stimulus onset were extracted from the continuous EEG data. The epochs did not include the onset of the US to preserve trials in the CS+ conditions that can be disproportionally affected by movement artifacts produced in anticipation of the US onset.

Artifact correction and rejection were carried out according to the Statistical Control of Artifacts in Dense Sensors Arrays (SCADS) procedure developed by Junghöfer et al. (2000). The SCADS approach computed a compound quality index for each participant, channel, and trial using three statistical parameters (Junghöfer et al., 2000) rectified (absolute) mean amplitude; standard deviation, maximum transient voltage change). The compound quality index was evaluated for each channel, and channels with high artifact content throughout the recording (globally bad channels) were flagged when the quality index exceeded 2.5 standard deviations above the median of the quality index. Globally bad channels were interpolated using a 2D spline interpolation method where all channels that survived artifact correction and rejection were used to estimate the values of the flagged channel. Channels at the trial level were evaluated using the same threshold for the quality index, and flagged channels were interpolated within that trial using the same spline method. Finally, trials were flagged for rejection if the compound quality index exceeded 1.25 times the median of the quality index. An average of 4% of channels in the acquisition phase were interpolated across participants (min: 0 channels; max: 15 channels), and an average of 91.60% of the total trials in the acquisition phase were retained for further analysis (tritan CS+ = 27.64; tritan CS-= 26.94; luminance CS+ = 27.86; luminance CS-= 27.48 trials on average out of 30).

The preprocessed and trial-averaged EEG data was prepared for the Hilbert transform to extract the time course of the ssVEP amplitude envelope for each condition (Figueira et al., 2022). To this end, the EEG data was averaged across conditions for individual subjects, average referenced, and baseline corrected to the average voltage of the 200 ms prior to stimulus onset. Next, the baseline corrected data was filtered using two 9^th^ order Butterworth filters with an upper cutoff frequency of 8 Hz and the other with a lower cutoff frequency of 7 Hz. The narrow filtering extracted the frequency that the gratings flickered at (7.5 Hz), so the results could be interpreted in a frequency band specific manner (Cohen, 2014). The filtered data was Hilbert transformed using the Matlab *hilbert.m* function to extract the time-varying estimates of phase and amplitude from the analytic signal over the entire epoch (−600 ms - 3800 ms), using the freqtag_hilbert.m function from the Matlab toolbox by Figueira et al. (2022). The Hilbert transform computes the analytic signal by doubling the positive-frequency Fourier coefficients, zeroing the negative-frequency coefficients, and taking the inverse of the Fourier transform of the doubled and zeroed Fourier coefficients (Cohen, 2014). In the time domain, the analytic signal is a phase-shifted version of the original narrowly bandpassed time series.

### Statistical analyses

US expectancy ratings were analyzed using paired t-tests. The expectancy ratings after the habituation phase were examined to verify that there were no stimulus driven differences in expectancy in the absence of the US. Moreover, expectancy ratings during acquisition were analyzed to validate the aversive conditioning design, and expectancy ratings for the extinction phase were included for ethical reasons.

The ssVEP responses were analyzed using a non-parametric Bayesian bootstrapped approach to estimate support for the hypothesis that the ssVEP amplitude at a given time point and sensor was larger for the CS+ stimuli compared to the CS-stimuli in the luminance and tritan conditions. To this end, the ssVEP envelope difference across stimulus types was calculated for each sensor and time point in the acquisition phase only. As suggested by Efron (2011), evidence supporting the hypothesis was quantified by bootstrapping the condition differences (2000 iterations) as well as a null distribution (calculated through permuting condition labels within subjects, 2000 iterations). Bootstrapped distributions were generated by randomly drawing a participant’s amplitude difference with replacement, for each iteration (see Ahumada et al. 2025 for a detailed description). The null distribution was generated by randomly drawing a participant’s amplitude difference after randomizing the conditions with replacement, for each iteration. Logarithmic Bayes factors (log10BF) were computed as the posterior odds that the differences were outside the null distribution, relative to uninformative (flat) prior odds of 1. Greater evidence to support the null hypothesis (BF01) would have suggested that there were no ssVEP amplitude differences between the CS+ and CS-stimuli. Bayes factors were interpreted using Jeffery’s scale (1961), where values ranging from .5 to 1 indicated substantial support, 1 to 1.5 indicated strong support, 1.5 to 2 indicated very strong support, and values greater than 2 indicated decisive support for our hypothesis. Furthermore, transitive Bayes factors were computed as the ratio of Bayes factors (interpreted using Jeffrey’s scale) across conditions to compare the magnitude of the tritan and luminance conditioning effects.

Finally, pupil dilation was analyzed using a Bayesian repeated-measures ANOVA to test the hypothesis that CS+ trials would evoke greater pupil re-dilation compared to CS-trials regardless of condition.

### Code accessibility

The custom preprocessing, postprocessing, and statistical analysis functions can be found at https://osf.io/yue78/.

## Results

### US Expectancy Ratings

Figure 2 shows the mean expectancy for the CS+ and CS-stimuli at each rating occasion (habituation, H; acquisition 1, A1; acquisition 2, A2; extinction, E) in the tritan and luminance conditions. The expectancy ratings were compared across stimuli at each rating occasion and for each condition using paired samples t-tests computed in JASP (JASP Team, 2024). US expectancy analysis revealed that that there were no significant differences in expectancy across stimulus type for the habituation rating occasion in the luminance (t(49) = 1.17, p = .250, *d* = 0.17) or tritan condition (t(49) = 1.59, p = .119, *d* = 0.23). However, for each acquisition rating occasion, the CS+ stimulus elicited significantly higher expectancy compared to the CS-stimulus for both the luminance (A1: t(49) = 7.11, p < .001, *d* = 1.01; A2: t(49) = 11.06, p < .001, *d* = 1.56) and tritan conditions (A1: t(49) = 6.63, p < .001, *d* = .94; A2: t(49) = 11.13, p < .001, *d* = 1.57). Moreover, expectancy ratings were significantly higher for CS+ stimulus compared to CS-stimulus for the extinction rating occasion in the luminance (t(49) = 7.67, p < .001, *d* = 1.08) and tritan conditions (t(49) = 7.94, p < .001, *d* = 1.12). Therefore, fear conditioning was successful across both experimental conditions.

**Figure 2.**
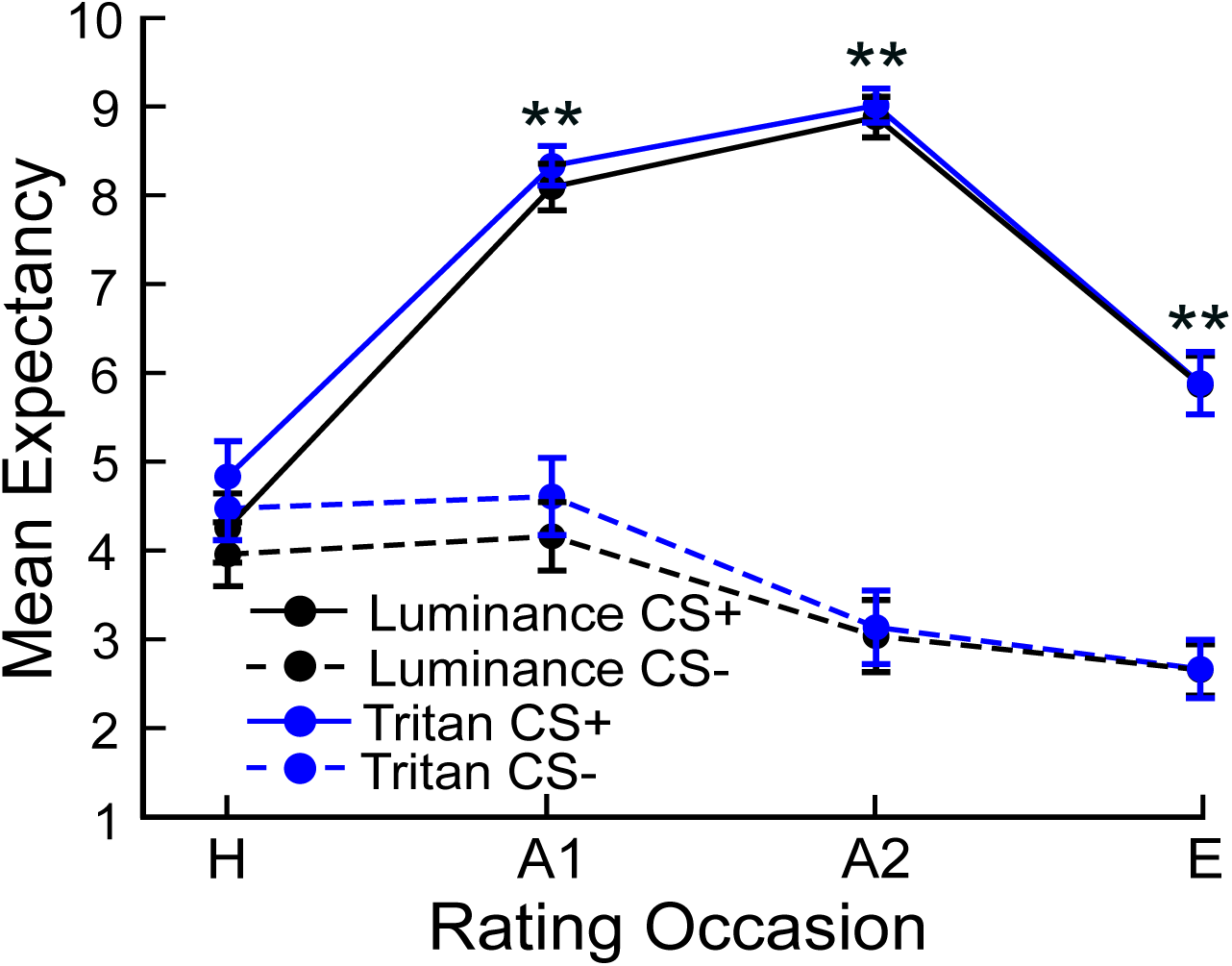
US expectancy ratings. Mean US expectancy ratings where 0 indicated low expectancy that the flickering grating was accompanied by a shock, 5 indicated uncertainty, and 10 indicated high expectancy that the flickering grating was accompanied by a shock. The habituation and extinction phases included one rating occasion (H and E), and the acquisition phase included two rating occasions (A1, 20 trials into acquisition; A2, 80 trials into acquisition). Paired t-tests indicated that US expectancy ratings for the CS+ stimuli were significantly higher compared to the ratings for the CS-stimuli during the acquisition and extinction rating occasions, regardless of condition.

### Steady-state Visual Evoked Potentials

Figure 3A shows the frequency spectrum of the visuocortical activity at sensor Oz (75) in the tritan CS+ condition across all participants in the acquisition phase. The spike in the frequency spectrum at 7.5 Hz indicates that the driving frequency was present in the spectrum. Furthermore, figure 3B shows the results of the Hilbert transform at sensor Oz (75) across participants for each trial type in each condition, indicating that the driving frequency elicited visuocortical amplification regardless of condition.

**Figure 3.**
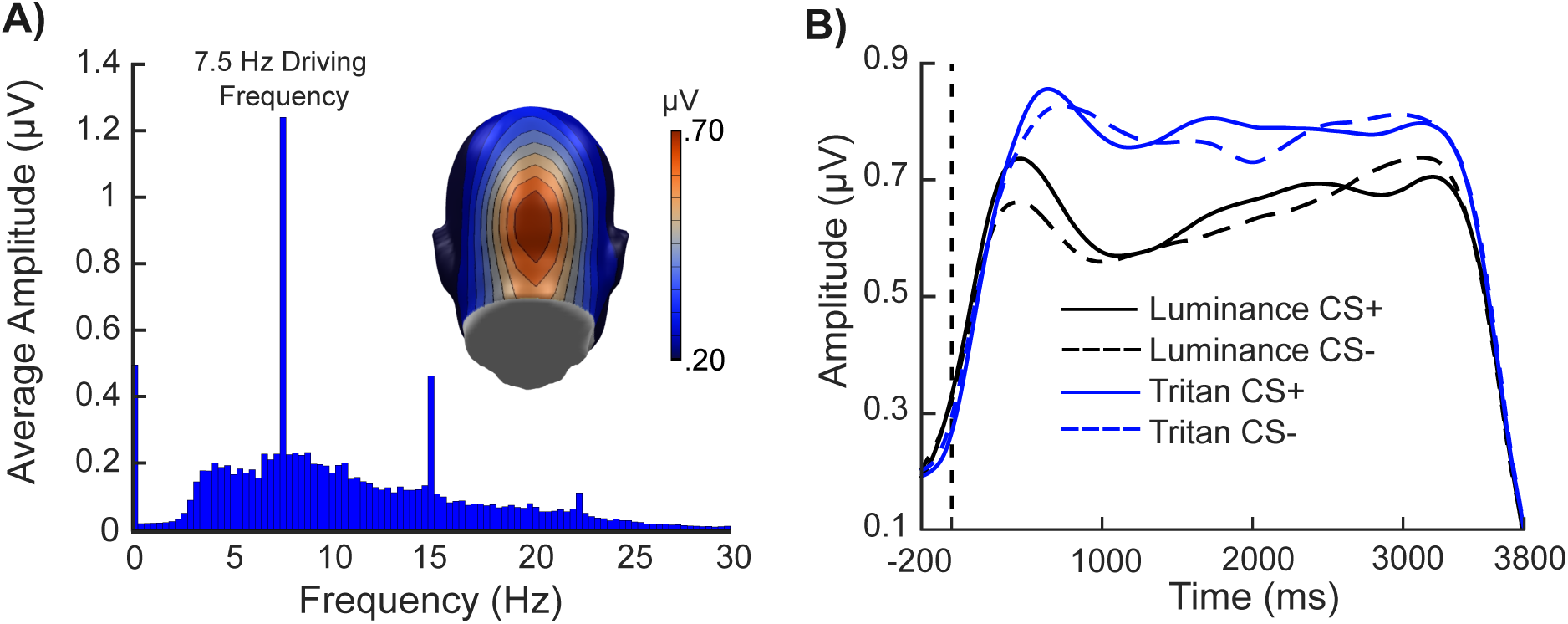
Grand mean (N = 50) ssVEP results at sensor Oz (75). The Fourier and Hilbert transformed ssVEP results. (A) The frequency spectrum of the ssVEP response to the tritan CS+ stimulus collapsed across participants at sensor Oz. The spike in the frequency spectrum at 7.5 Hz indicates the presence of the driving frequency. (B) The results of the Hilbert transform at sensor Oz for all conditions and stimulus types collapsed across participants.

Figure 4 shows the results of the non-parametric Bayesian bootstrapped approach used to estimate support for the hypothesis that the ssVEP amplitude at a given time point and sensor was larger for the CS+ stimuli compared to the CS-stimuli in each condition. For the luminance condition, an early conditioning effect (0 ms - 1000 ms) showed decisive support for our hypothesis (averaged log10BF > 2, peak log10BF = 2.07 at 620 ms) in a right-lateralized occipital cluster of sensors (75, 83 (O2), 84, 89, and 90; luminance O2 cluster). Similarly, an early conditioning effect was observed in the tritan condition, but in a left-lateralized occipital cluster of sensors (65, 66, 69, 70 (O1), and 75; tritan O1 cluster) that showed decisive support for our hypothesis (averaged log10BF > 2, peak log10BF = 2.22 at 576 ms). Furthermore, a late conditioning effect (1500 ms - 2500 ms) in the tritan condition was observed in a cluster of several anterior sensors over the right and left hemispheres (29, 30, 35, 36 (C3), 80, 87, 104 (C4), and 105; tritan anterior cluster) and showed decisive support for our hypothesis (averaged log10BF > 2, peak log10BF = 2.10 at 2206 ms).

**Figure 4.**
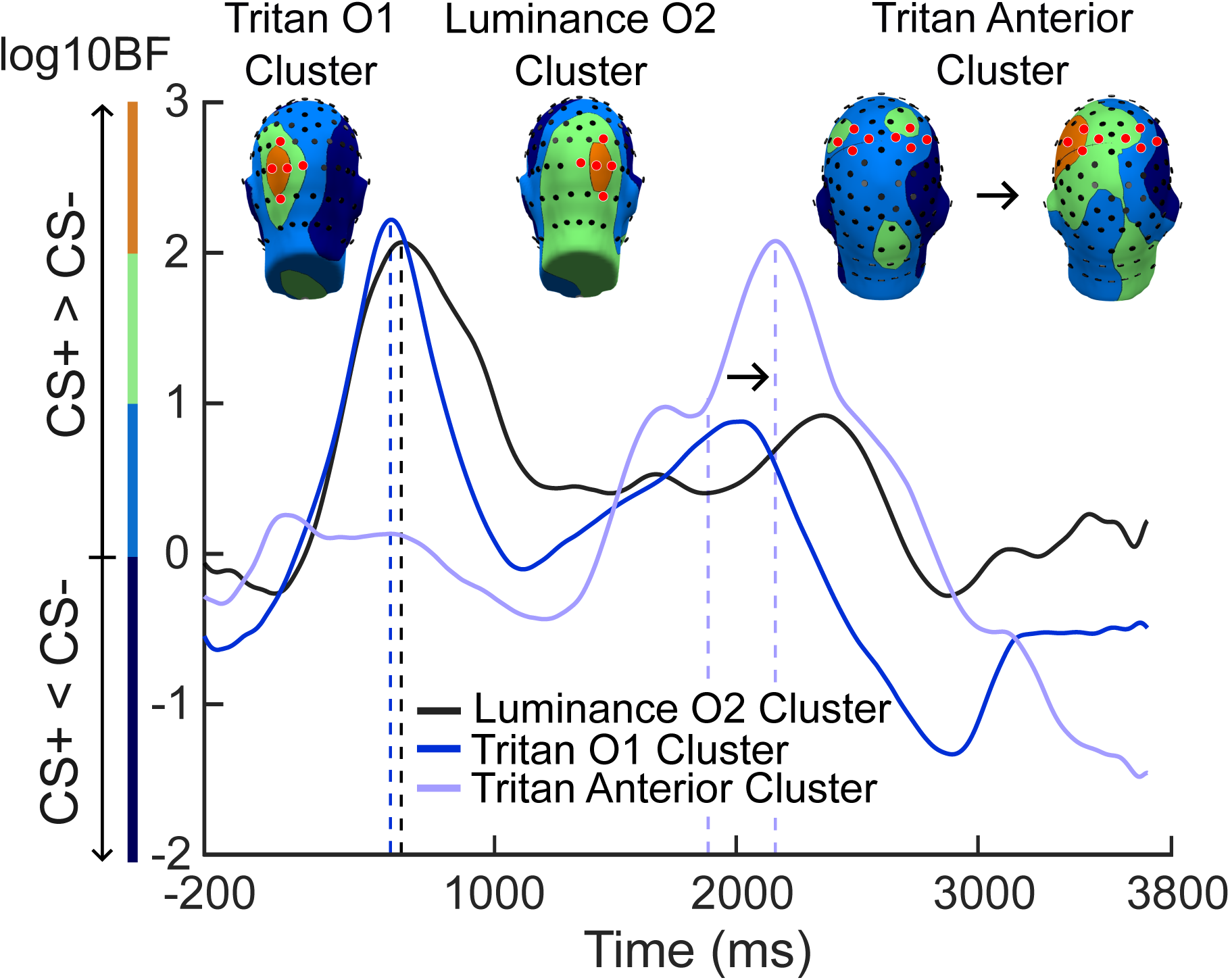
Effects of aversive conditioning. The condition level averaged log10BF values of the three clusters (sensors indicated in red) of electrodes that showed decisive support for our hypothesis that the ssVEP amplitude at a given time point and sensor was larger for CS+ stimuli compared to CS-stimuli, regardless of condition. Log10BF values were interpreted using Jeffery’s scale (1961) where values ranging from .5 to 1 indicated substantial support, 1 to 1.5 indicated strong support, 1.5 to 2 indicated very strong support, and values greater than 2 indicated decisive support for our hypothesis. Cluster O2 included sensors 75, 83 (O2), 84, 89, and 90; cluster O1 included sensors 65, 66, 69, 70 (O1), and 75; and the anterior cluster included sensors 29, 30, 35, 36 (C3), 80, 87, 104 (C4), and 105. The occipital topographies were plotted at the time point where the occipital clusters reached the peak logBF10 value (tritan O1 cluster blue dashed line, peak log10BF = 2.22 at 576 ms; luminance O2 cluster black dashed line, peak log10BF = 2.07 at 620 ms). The anterior topographies were plotted at the time point where the bilateral effect emerged and also reached the peak logBF10 value (tritan anterior cluster, first purple dashed line, log10BF = 1.84 at 1950 ms; second purple dashed line, peak log10BF = 2.10 at 2206).

Figure 5 shows the Bayesian bootstrapped z-transformed conditioning effect and null distributions for sensors and time points corresponding to the clusters identified in figure 4. The early luminance distributions were computed for sensor O2 at 620 ms, the early tritan distributions were computed for sensor O1 at 576 ms, and the late tritan distributions were computed for sensor C4 at 1950 ms and sensor C3 at 2206 ms. The early luminance distributions at sensor O2 showed less overlap between the null and effect distributions, compared to the early tritan distributions at sensor O1. Furthermore, the late tritan distributions at sensor C3 showed less overlap between the null and effect distributions, compared to the late tritan distributions at sensor C4.

**Figure 5.**
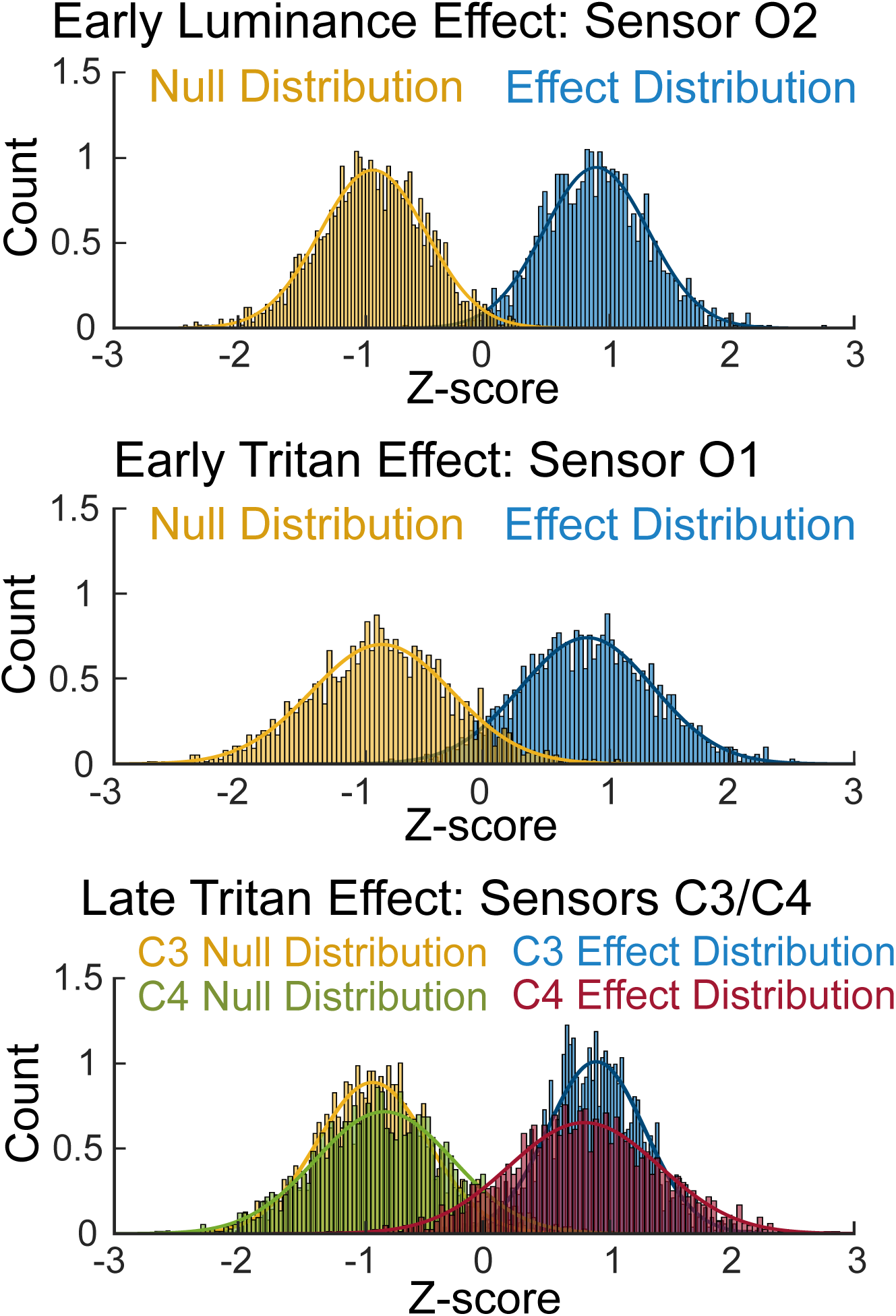
Bayesian bootstrapped z-transformed distributions for the early and late conditioning effects. Density plots of Bayesian bootstrapped z-transformed null and conditioning effect distributions at sensors O1, O2, C3, and C4. Distributions for the early and late conditioning effects were computed at the sensor and time point corresponding to the cluster identified in figure 4: early luminance effect (sensor O2 at 620 ms), early tritan effect (sensor O1 at 576 ms), and late tritan effect (sensor C4 at 1950 ms; sensor C3 at 2206 ms). Distributions included 2000 draws. Note that the distributions were z-transformed together to facilitate distribution fitting, so the null-distributions are not centered around zero.

Figure 6 shows the transitive Bayes factors (logBF10) mapped across all sensors for the early and late time windows that were identified in figure 4. Transitive Bayes factors were computed as the ratio of Bayes factors in the tritan condition to the Bayes factors in the luminance condition. Therefore, negative transitive Bayes factors favored the luminance conditioning effect over the tritan conditioning effect, and positive transitive Bayes factors favored the tritan conditioning effect over the luminance conditioning effect at a given time point and sensor. Transitive Bayes factors showed decisive evidence of a larger early luminance conditioning effect compared to the early tritan conditioning effect (0 ms - 1000 ms) in a considerable number of central and right-lateralized occipital sensors (log10BF > 2; peak log10BF = −3.13, sensor Pz; peak log10BF = −3.06, sensor 90). Furthermore, the early tritan conditioning effect was larger than the early luminance conditioning effect in only a small number of non-adjacent left-lateralized occipital sensors 47, 52, and 65 (log10BF > 2; peak log10BF = 2.40, sensor 65). Moreover, transitive Bayes factors showed decisive support for the prominence of the late tritan conditioning effect (1500 ms - 2500 ms) in a substantial number of anterior bilateral sensors (log10BF> 2; peak log10BF = 3.37, sensor C3, left hemisphere; peak log10BF = 2.83, sensor C4, right hemisphere).

**Figure 6.**
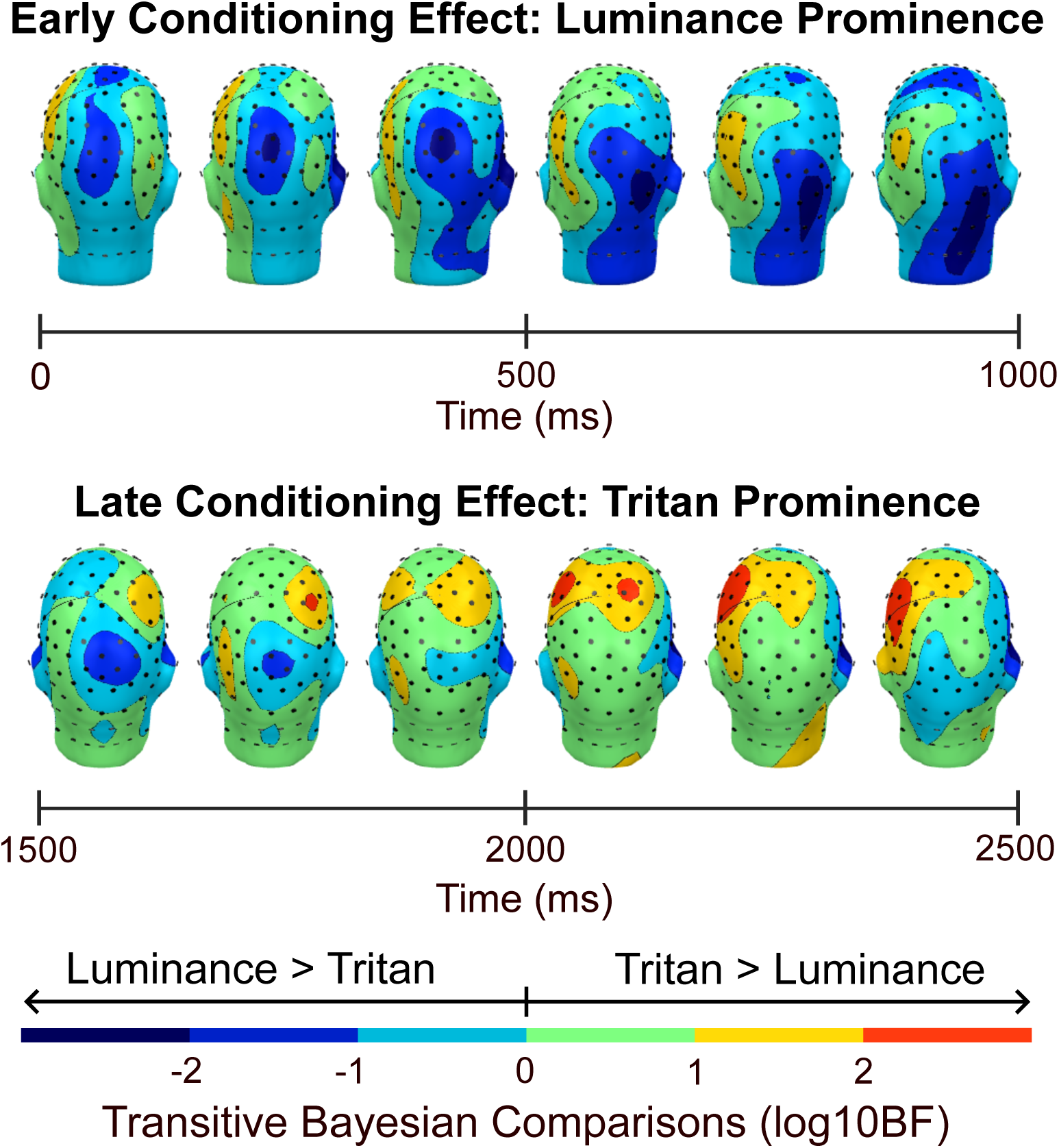
Transitive log10BF topographic map. Topographic map of transitive Bayes factors computed as the ratio of the Bayes factors of the tritan conditioning effect to the Bayes factors of the luminance conditioning effect. Negative transitive Bayes factors indicated that the luminance conditioning effect was larger than the tritan conditioning effect, and positive transitive Bayes factors indicated that the tritan conditioning effect was larger that the luminance conditioning effect. Transitive Bayes factor values were interpreted using Jeffery’s scale (1961) where absolute log10BF values ranging from .5 to 1 indicated substantial support, 1 to 1.5 indicated strong support, 1.5 to 2 indicated very strong support, and values greater than 2 indicated decisive support for our hypothesis. An early occipital effect (0 ms - 1000 ms) showed decisive support favoring the luminance conditioning effect over the tritan conditioning effect, indicating the prominence of the early luminance conditioning effect. Additionally, a late anterior effect (1500 ms - 2500 ms) showed decisive support favoring the tritan conditioning effect and demonstrated the prominence of the late tritan conditioning effect.

### Pupil

Figure 7 shows the results of the 2 (Condition) X 2 (Stimulus type) Bayesian repeated measures ANOVA computed in JASP (JASP Team, 2024) that examined the effect of condition (luminance and tritan) and stimulus type (CS+ and CS-) on pupil diameter (mm) over the last two seconds of the epoch (1800 ms - 3800 ms). The results indicated decisive support (log10BF = 32.98) for the model that included main effects of Condition and Stimulus type, and there was no evidence of an interaction. Bayesian post-hoc tests showed decisive evidence of differences in pupil diameter across conditions (log10BF = 63.20) and stimulus types (log10BF = 2.91). Pupil diameter, regardless of stimulus type, was smaller in the luminance condition (CS+, M = −0.60 mm; CS-, M = −0.66 mm) compared to the tritan condition (CS+, M = 0.22 mm; CS-, M = 0.16 mm), but the CS+ stimuli elicited greater pupil re-dilation compared to the CS-stimuli regardless of condition.

**Figure 7.**
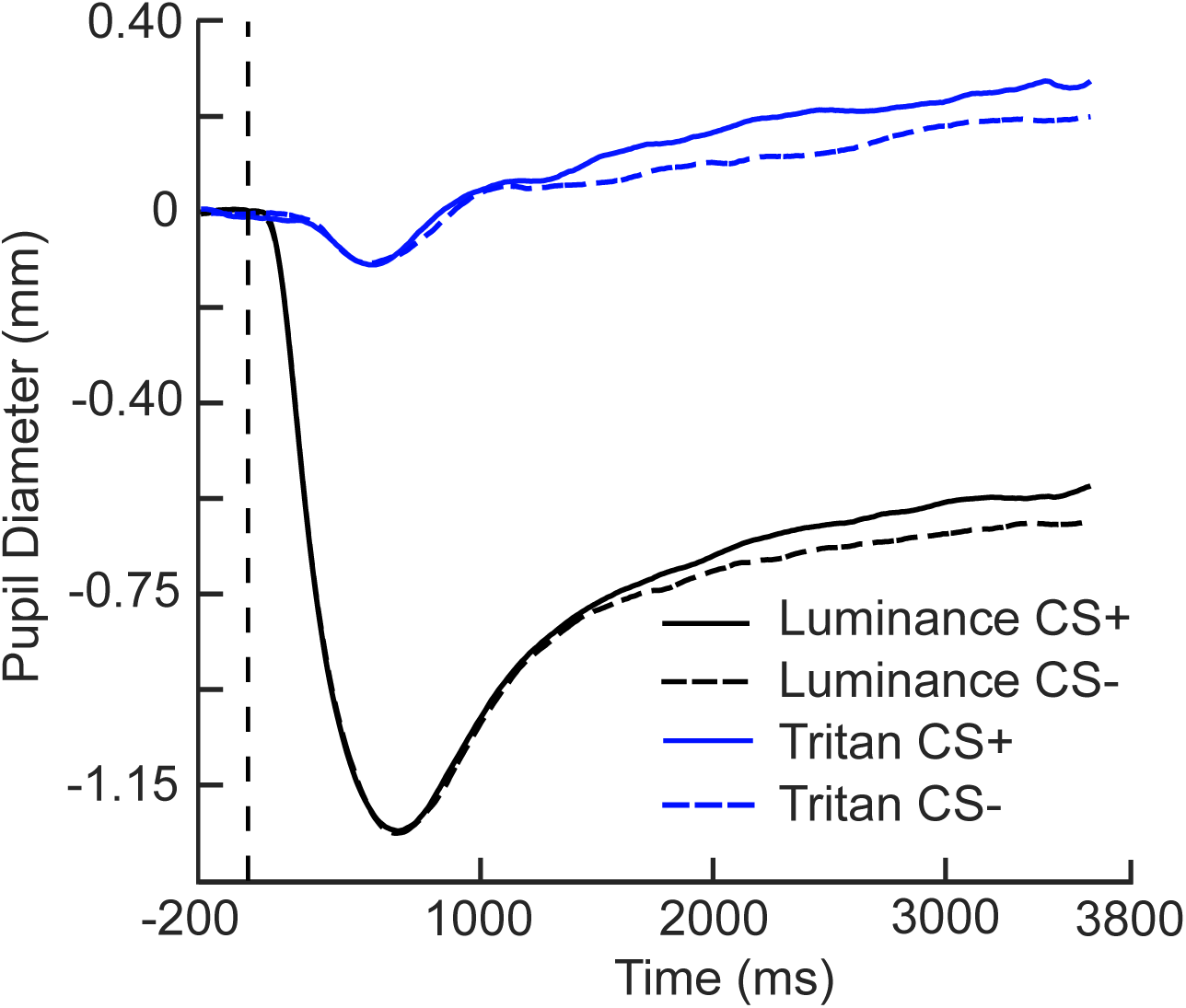
Grand mean (N = 50) pupil diameter. Changes in pupil diameter over time for each condition and stimulus type collapsed across all participants. The measurement window included in the 2 (Condition) X 2 (Stimulus) Bayesian repeated measures ANOVA was 2800 ms to 3800 ms to assess differences in pupil re-dilation.

## Discussion

The present study investigated the contribution of the koniocellular visual pathway to aversive learning in human visual cortex using steady-state visual evoked potentials to examine selective electrocortical amplification of threat cues. Tritan and luminance channel biasing stimuli were used to convey visual information to the koniocellular visual pathway and luminance channels, respectively. We tested the hypothesis that aversive conditioning would enhance visuocortical activation to threat-related stimuli in the tritan condition, paralleling established findings for stimuli in which the full luminance information is available. As expected, the CS+ stimuli evoked increased ssVEP responses in both the tritan and luminance conditions, compared to the CS-stimuli. Additionally, transitive Bayes factors were computed to evaluate the magnitude of the conditioning effects across the tritan and luminance conditions. Our results showed decisive support that favored the early luminance conditioning effect over the early tritan conditioning effect, indicating that the early luminance conditioning effect was larger than the early tritan conditioning effect in several central and right lateralized occipital sensors early into the trial (0 ms - 1000 ms). Furthermore, decisive evidence indicated the presence of a prominent late tritan conditioning effect in several bilateral anterior sensors at around 1500 ms to 2500 ms post-stimulus onset.

In line with our first hypothesis, tritan stimuli elicited robust ssVEP responses overall, extending previous findings of visuocortical amplification of S-cone biasing stimuli (Cottaris and De Valois, 1998; Mullen et al., 2007, 2008). Consistent with our second hypothesis, we observed greater visuocortical activation in response to the CS+ stimuli, compared to the CS-stimuli in both the tritan and luminance conditions. Moreover, decisive support was shown for the conditioning effects in each condition (logBF10 > 2), suggesting that activity along the koniocellular pathway participates in the representation of conditioned threat in humans.

To further elucidate the koniocellular contribution to aversive learning, transitive Bayes factors were computed to compare the magnitude of the conditioning effects across conditions. Transitive Bayes factors showed decisive evidence favoring the early tritan conditioning effect in a small number of non-adjacent left-lateralized occipital sensors. However, transitive Bayes factors also revealed decisive evidence indicating the prominence of the early luminance conditioning effect over the early tritan conditioning effect in a substantial number of adjacent central and right-lateralized occipital sensors. Our findings align with aversive conditioning studies that used luminance channel biasing stimuli and observed robust visuocortical amplification of threat-related stimuli compared to safe stimuli over central and right-lateralized occipital sensors (Friedl and Keil, 2021; Stegmann et al., 2023). Moreover, the topographies of the early conditioning effects were consistent with the notion that early visual areas contribute to the generation of ssVEP responses (Di Russo et al., 2006).

Early visual cortex contributes to the selective amplification of aversive stimuli through feedforward projections from V1, local inhibitory circuits, and feedback projections from high-order visual areas (Angelucci et al., 2002; Shapley et al., 2003, 2007; Keil et al., 2007). Therefore, the difference in the magnitude of the early conditioning effects could be attributed to the specific neurophysiological connections associated with each channel. The input layer of V1 (Layer 4C) receives substantial excitatory geniculo-cortical innervation, including magnocellular and parvocellular geniculo-cortical projections (Hubel and Wiesel, 1972; Livingstone and Hubel, 1984; Callaway, 1998; Garcia-Marin et al., 2019). Conversely, the koniocellular pathway exhibits distributed geniculo-cortical projections to the output layers of V1, including layer 1 and the cytochrome oxidase blobs in layer 2/3 in primates (Livingstone and Hubel, 1984; Hendry and Yoshioka, 1994; Hendry and Reid, 2000).

It has been hypothesized that the laminar layers of V1 are functionally distinct, and several studies have modeled how the functional property of orientation selectivity emerges in V1 across the laminar layers (Sompolinsky and Shapley, 1997; Shapley et al., 2003; Gur et al., 2005). Ringach and colleagues (2002) provided evidence of neurons in V1 that display diverse orientation selectivity and contribute to broadly distributed orientation selectivity across the layers of V1 in primates. Moreover, Gur and colleagues (2005) found that the distribution of orientation tuning in V1 in primates is not uniform, and differences in tuning could be attributed to the variability of excitatory and inhibitory mechanisms in each layer of V1. For instance, local inhibition is thought to sharpen the broadly tuned excitatory projections from the LGN and may facilitate the first stage of steady-state orientation specific responses observed over early visual areas (Shapley et al., 2003, 2007). Moreover, feedback projections from higher-order visual areas are also thought to play a role in modulating the magnitude of the ssVEP by conveying information about arousal to early visual areas and displaying a pattern of retinotopic alignment with feedforward projections from V1 in primates (Angelucci et al., 2002; Buffalo et al., 2010; Briggs, 2020; Li and Keil, 2023). Therefore, differences in high-order feedback projections and local inhibitory interactions that shape the ssVEP responses could contribute to the prominence of the luminance-driven early conditioning effect over the early tritan conditioning effect.

Evidence of a late anterior conditioning effect was observed in the tritan condition at around 1500 ms to 2500 ms and showed decisive support for the hypothesis that the ssVEP responses to the tritan CS+ stimuli were greater than the responses to the tritan CS-stimuli. Moreover, transitive Bayes factors showed decisive support indicating the prominence of a late tritan conditioning effect in several bilateral anterior sensors. Although ssVEP responses are primarily generated in early visual cortex and tend to emerge maximally over occipital sensors, recent work has established that area MT is also involved in ssVEP generation (Di Russo et al., 2006; Petro et al., 2017). For instance, Boylan and colleagues (2023) observed that attention-modulated ssVEP responses to luminance gratings were associated with BOLD activation in area MT. Additionally, functional imaging evidence in humans showed that S-cone biasing stimuli elicit activity in area MT (Wandell et al., 1999; Mullen et al., 2008). Therefore, future investigations using neuroimaging methods with better spatial resolution are needed to address the potential involvement of extrastriate cortex in the late visuocortical amplification of threat signals in the tritan condition.

The present study extends our knowledge of the properties of the koniocellular visual pathway, providing evidence of the tritan contribution to aversive learning in humans. Moreover, we replicated previous findings showing enhanced visuocortical activation in response to tritan stimuli, and our findings suggest that both the koniocellular pathway and luminance-sensitive channels play a role in the visuocortical enhancement of threat signals in human visual cortex.

## Acknowledgments

This research was supported by grant R01MH125615 from the National Institutes of Health.

## Notes

The authors declare no conflicts of interest.

https://osf.io/yue78/

### Competing Interest Statement

The authors have declared no competing interest.

### Summary of Updates

An update to the title and minor manuscript revisions were completed to address editorial comments and to improve the clarity of the methodology. Visual stimuli were measured using a Gossen MavoSpot Luminance meter.

